# Genome-wide Analysis In Response to N and C Identifies New Regulators for root AtNRT2 Transporters

**DOI:** 10.1101/822197

**Authors:** Sandrine Ruffel, Valentin Chaput, Jonathan Przybyla-Toscano, Ian Fayos, Catalina Ibarra, Tomas Moyano, Cécile Fizames, Pascal Tillard, Jose Antonio O’Brien, Rodrigo A. Gutiérrez, Alain Gojon, Laurence Lejay

## Abstract

In *Arabidopsis thaliana*, the High-Affinity Transport System (HATS) for root NO_3_^-^ uptake depends mainly on four NRT2 transporters, namely NRT2.1, NRT2.2, NRT2.4 and NRT2.5. The HATS is the target of many regulations to coordinate nitrogen (N) acquisition with the N status of the plant and with carbon (C) assimilation through photosynthesis. At the molecular level, C and N signaling pathways have been shown to control gene expression of the *NRT2* transporters. Although several regulators of these transporters have been identified in response to either N or C signals, the response of *NRT2* genes expression to the interaction of these signals has never been specifically investigated and the underlying molecular mechanisms remain largely unknown. To address this question we used an original systems biology approach to model a regulatory gene network targeting *NRT2.1, NRT2.2, NRT2.4* and *NRT2.5* in response to N/C signals. Our systems analysis of the data highlighted the potential role of three putative transcription factors, TGA3, MYC1 and bHLH093. Functional analysis of mutants combined with yeast one hybrid experiments confirmed that all 3 transcription factors are regulators of *NRT2.4* or *NRT2.5* in response to N or C signals.

**One sentence summary:** Identification of three transcription factors involved in the regulation of NRT2s transporters using a systems biology approach and NRT2.1 as target gene in response to combinations of N/C treatments

## Introduction

As all living organisms, plants must integrate internal and external signals to adapt to fluctuating environmental conditions. This is particularly the case concerning mineral nutrition, because most nutrients display dramatic changes in external availability, whereas their internal concentrations must be kept within a limited range to be compatible with physiological processes. Accordingly, root nutrient uptake systems are finely tuned by regulatory mechanisms activated by local signaling of external nutrient availability and systemic signaling of the nutrient status of the whole plant (Schachtman and Shin, 2007). Furthermore, acquisition of the various nutrients has to be coordinated to remain consistent with the global chemical composition of plant tissues and with the fact that most nutrients contribute to the synthesis of biomolecules with a relatively strict elemental stoichiometry (e.g., C, N and S for amino acids). Therefore, the signaling pathways that are specific for the different nutrients must interact to ensure this coordination. Although coordinated regulation of uptake systems for different nutrients have been clearly demonstrated at the physiological level, the underlying molecular mechanisms remain largely obscure (Schachtman and Shin, 2007). The cross-talks between N and C signaling mechanisms are certainly those that have been most often investigated (Coruzzi and Zhou, 2001; Nunes-Nesi et al., 2010; Ruffel et al., 2014), first because N and C are the two mineral nutrients plants require in largest quantities, and also because they connect two key functions of plants as autotrophic organisms, *i.e.*, photosynthesis and assimilation of inorganic nitrogen. Moreover, the importance of N/C signaling interaction is dramatically illustrated by the fact that most N-responsive genes in *Arabidopsis* are actually regulated by C/N interaction (Gutierrez et al., 2007).

The nitrogen nutrition of most herbaceous plants relies on the uptake of nitrate (NO_3_^-^), which is ensured in root cells by two classes of transport systems. The High-Affinity Transport System (HATS) is predominant in the low range of NO_3_^-^ concentrations (up to ∼ca 1 mM), whereas the Low-Affinity Transport System (LATS) makes an increasing contribution to total NO_3_^-^ uptake with increasing external NO_3_^-^ concentration (Crawford and Glass, 1998). In all species investigated to date, genes encoding the various transporter proteins involved in either HATS or LATS have mostly been identified in the *NRT2* and *NPF* (formerly *NRT1/PTR*) families, respectively (Nacry et al., 2013; O’Brien et al., 2016). The respective roles of HATS and LATS in the total NO_3_^-^ acquisition by the plant are still a matter of debate. However, field studies suggest that even in agricultural conditions, the HATS has a major contribution over the whole developmental cycle (Malagoli et al., 2004; Garnett et al., 2013). Both the structure and regulation of the HATS have been extensively studied in *Arabidopsis thaliana*. In this species, almost all the HATS activity depends on four NRT2 transporters, namely NRT2.1, NRT2.2, NRT2.4 and NRT2.5 (Filleur et al., 2001; Kiba et al., 2012; Lezhneva et al., 2014), which all require an interaction with the NAR2.1 protein to be active in NO_3_^-^ transport (Kotur et al., 2012). Under most conditions, NRT2.1 is the main contributor to the HATS (Cerezo et al., 2001; Filleur et al., 2001). However, NRT2.4 and NRT2.5 display a very high-affinity for NO_3_^-^ and are important for taking up this nutrient when present at very low concentration (<50 µM) in the soil solution (Kiba et al., 2012; Lezhneva et al., 2014). Furthermore, unlike NRT2.1 and NRT2.4, NRT2.5 does not require the presence of NO_3_^-^ to be expressed, and is therefore considered crucial for ensuring the initial uptake of NO_3_^-^ as soon as it appears in the external medium (Kotur and Glass, 2015).

Most interestingly, the HATS has been shown to be the target of almost all regulations governing root NO_3_^-^ acquisition in *Arabidopsi*s (Nacry et al., 2013), and this is associated with control of *NRT2.1, NRT2.2, NRT2.4* and *NRT2.5* expression at the mRNA level. In particular, previous reports have shown that *NRT2.1* is induced both by N starvation (Lejay et al., 1999; Cerezo et al., 2001; Gansel et al., 2001), and by light and sugars, indicating coordination with photosynthesis (Lejay et al., 1999; Lejay et al., 2003). This makes *NRT2.1* a very relevant model gene for investigating the interaction between N and C signalling networks in roots. This also holds true for *NRT2.4* (Lejay et al., 2008; Kiba et al., 2012), but not for *NRT2.5*, which until now has only been reported to be up-regulated by N starvation (Lezhneva et al., 2014). For these reasons, and also due to its high functional importance as the main component of the HATS, *NRT2.1* has been extensively investigated to unravel its regulatory mechanisms. Accordingly, a quite significant number of genes have been found to encode regulators of *NRT2.1* expression, such as *LBD37-39* (Rubin et al., 2009), *TGA1* and *TGA4* (Alvarez et al., 2014), *NLP6* and *NLP7* (Marchive et al., 2013; Guan et al., 2017), *NRG2* (Xu et al., 2016), *BT1-2* (Araus et al., 2016), *NRT1.1* (Munos et al., 2004), *CIPK8* (Hu et al., 2009), *HNI9/IWS1* (Widiez et al., 2011) and *HY5* (Chen et al., 2016). Most of these genes contribute to the regulation of *NRT2.1* expression in response to changes in N provision. The only exception is *HY5*, which encodes a transcription factor reported to ensure long-distance signalling of the stimulation of *NRT2.1* expression in roots by illumination of the shoot. Strikingly, none of the above regulators were shown to be involved in the cross-talk between N and C signalling pathways. Even more surprising, the response of *NRT2.1* expression itself (as well as those of the other *NRT2s*) to the interaction of N and C signals was not specifically investigated. As a consequence, the molecular mechanisms responsible for the coordinated regulation of the NO_3_^-^ HATS by N and C status of the plant are unknown. Our study aimed at filling this gap. Therefore, using *NRT2.1* as a marker gene to identify relevant combinations of N/C treatments, we developed a systems biology approach based on genome-wide transcriptome analysis in roots of *Arabidopsis* plants to model a regulatory gene network targeting *NRT2.1, NRT2.2, NRT2.4* and *NRT2.5* in response to N/C signals. This highlighted the potential role of three putative transcription factors, TGA3, MYC1 and bHLH093 in controlling the expression of these transporter genes. Functional analysis of loss-of-function mutants confirmed that all 3 transcription factors are regulators of *NRT2.4* or *NRT2.5* in response to N or C signals. Furthermore, yeast one hybrid experiments confirmed that at least TGA3 and MYC1 are able to bind *NRT2.4* and *NRT2.5* promoters.

## Results

### Regulation of root nitrate transporters by interaction between nitrogen and light provision

We wished to determine whether induction of *NRT2.1* by N starvation is dependent on light, and conversely if *NRT2.1* induction by light is dependent on the availability of NO_3_^-^ (Figure 1A and Figure 1B). In order to reveal possible interactions between C and N signalling pathways for the regulation of *NRT2.1*, we performed two different sets of experiments. In the first set of experiments, plants were starved for N for up to 72h either in the dark or at three different light intensities, 50 μmol m^-2^s^-1^ (LL), 250 μmol m^-2^s^-1^ (IL) and 800 μ mol m^-2^s^-1^ (HL) (Figure 1A). In the second set of experiments, plants were treated with 10mM NO_3_^-^, 1mM NO_3_^-^ or no N and transferred during 8h from the dark to HL conditions (Figure 1B).

**Figure 1.**
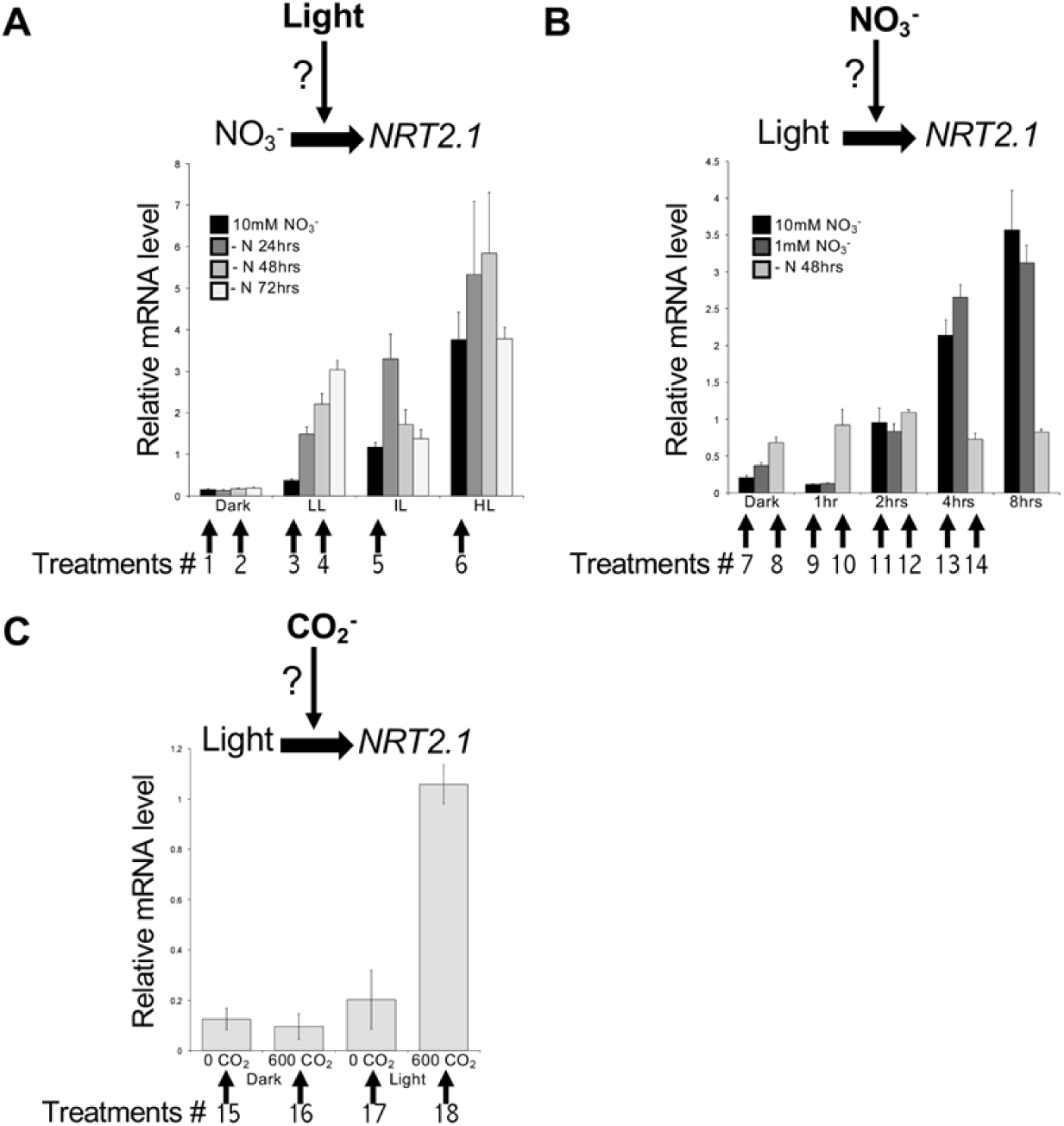
Interaction between Nitrogen and Light/Carbon provision modulates *NRT2.1* mRNA accumulation in roots. **A.** Different light regimes modulate *NRT2* regulation in roots of plants experiencing from high NO_3_^-^ provision (10 mM) to N deprivation (-N). The light regimes encompass dark, low light intensity (50 μmol m^-2^ s^-1^; LL), intermediate light intensity (250 μmol m^-2^ s^-1^; IL) and high light intensity (800 μmol m^-2^ s^-1^; HL). Plants were supplied with NO_3_^-^ 10 mM one week ahead the experiment and acclimated for 24 hours in the different light regimes before applying the N deprivation for 24, 48 or 72 hr. **B.** Different N provisions modulate *NRT2* regulation in roots of plants experiencing a dark to light transition. The N provisions encompass 10mM NO_3_^-^, 1mM NO_3_^-^ (for 72 hr) and N deprivation for 48 hr (-N). Plants are kept in the dark (*i.e.*, 40hr) before transition to high light intensity (800 μmol m^-2^ s^-1^; HL) and roots are collected at time 0 (Dark) and 1, 2, 4 and 8 hr after light transition. **C.** Regulation of *NRT2* by photosynthesis activity. Plants are grown in regular NO_3_^-^ regime (1mM) and intermediate light intensity until they are transferred for 4 hr in a CO_2_-deprived atmosphere (0ppm) or in high CO_2_-supplied atmosphere (600ppm), either in the dark or in the light. In these 3 experimental conditions, roots have been collected to assess *NRT2.1* mRNA accumulation by RT-QPCR (relative accumulation to *Clathrin* housekeeping gene). Expression pattern of *NRT2.1* across the 35 conditions tested (16 in A, 15 in B and 4 in C) has driven the choice of 18 conditions to investigate gene reprogramming associated to the regulation of NO_3_^-^ transport. These 18 conditions are indicated with arrows and numbers on the x-axis of the 3 *NRT2.1* bar graphs (Each arrow corresponds to one condition with 2 independent biological repeats constituted of a pool of approx. 10 plants each).

In LL and IL conditions, *NRT2.1* expression was, as expected, induced when plants were starved for N even if both the kinetic and the level of induction were different depending on light intensity (Figure 1A). When plants were kept in the dark, *NRT2.1* expression was not induced by N starvation but it remained very low both on 10mM NO_3_^-^ and on N free solution. More surprisingly, the induction of *NRT2.1* expression by N starvation was also almost completely abolished when plants were treated in HL conditions. However, under HL *NRT2.1* mRNA levels were always high, even under repressive conditions such as 10mM NO_3_^-^. This unexpected result is specific of *NRT2.1* since *NRT2.2, NRT2.4* and *NRT2.5*, known to be also induced by N starvation in roots (Li et al., 2007; Kiba et al., 2012; Lezhneva et al., 2014), are still regulated by N starvation in HL (Supplemental Figure 1A). However, just like *NRT2.1, NRT2.2, NRT2.4* and *NRT2.*5 were not regulated by N starvation in the absence of light. These data confirm the need of light for the regulation by N starvation of root NO_3_^-^ transporters. It also suggests that the mechanisms involved in *NRT2.1* regulation by N starvation are somewhat different from the mechanisms involved in the regulation of *NRT2.2, NRT2.4* and *NRT2.5*.

The second set of experiments confirmed the strong interaction between C/N signals as it revealed that the level of N nutrition affects the regulation of *NRT2.1* expression by light (Figure 1B). Indeed, when plants were starved for N during 48h, *NRT2.1* expression was much less induced by light as compared to plants grown on 10 or 1mM NO_3_^-^ (Figure 1B). Among other root NO_3_^-^ transporters, only *NRT2.2* and *NRT2.4* were induced by light and their level of induction seemed to be also dependent on N nutrition (Supplemental Figure 1B). However, in contrast to *NRT2.1*, the level of expression of both *NRT2.2* and *NRT2.4* was high when plants were starved for N and low when plants were grown on 1 or 10mM NO_3_^-^ (Supplemental Figure 1B). For *NRT2.4*, it confirms that this transporter is more sensitive to high N repression than *NRT2.1* (Kiba et al., 2012). The same result was obtained for *NRT2.5*, whose expression is barely detectable on either 10mM or 1mM NO_3_^-^ (Supplemental Figure 1B). However, concerning regulation by light, even when *NRT2.5* expression was high in N starved plants, light did not induce but rather seemed to repress *NRT2.5* mRNA accumulation after 8h in the light (Supplemental Figure 1B).

In a previous study, we showed that expression of *NRT2.1* and *NRT2.4* is induced by light only in the presence of CO_2_ in the atmosphere, suggesting that light regulation of these genes corresponds to a control exerted by photosynthesis (Lejay et al., 2008). As in the rest of our study we used micro-array experiments to look for genes involved in the regulation of root NO_3_^-^ transporters by photosynthesis, it was important for us to be able to discriminate between genes regulated by light itself or by photosynthesis. To do so, we performed a third set of experiments where plants were transferred from dark to light for 4h in an atmosphere containing 0 or 600ppm CO_2_. The results confirmed (i) that both *NRT2.1* and *NRT2.4* are only induced by light in the presence of CO_2_ and (ii) that *NRT2.5* is not induced by light or photosynthesis as suggested by our previous experiment (Figure 1C and Supplemental Figure 1C).

### Gene network for the regulation of root nitrate transporters by light and N starvation

The experiments performed above allowed us to reveal interesting interactions between C and N regulation of root NRT2 NO_3_^-^ transporters. We took advantage of this experimental design to develop a systems biology approach aiming at inferring a gene regulatory network underlying the interactions between N and C signals in the regulation of root NO_3_^-^ transporters. Due to the central position of *NRT2.1* as regulatory target affecting N acquisition, we used it as a focus gene around which to find associated gene networks.

We performed Affymetrix microarrays on selected combinations of light and N treatments, which were found discriminant for regulation of *NRT2.1*. Altogether, we chose 18 treatments labelled with numbered arrows in Figure 1. These 18 treatments correspond to 3 sets of conditions representative of (i) the light-dependent induction of *NRT2.1* expression in response to N starvation, (ii) the light induction of *NRT2.1* on 10mM NO_3_^-^ and (iii) the specific regulation of *NRT2.1* by photosynthesis and not by light itself. For each treatment, 2 independent biological replicates were generated and used for Affymetrix ATH1 microarray hybridization.

Regulation of *NRT2.1, NRT2.2, NRT2.4* and *NRT2.5* gene expression in response to N starvation and light/photosynthesis was similar on microarrays as compared to the results obtained by quantitative PCR in Figure 1 and Supplemental Figure 1 (Supplemental Figure 2). These results also confirmed that these four NO_3_^-^ transporters are the main *NRT2*s expressed in roots. *NRT2.3, NRT2.6* and *NRT2.7* showed very low expression levels on the microarrays under our experimental conditions. It is also noteworthy that *NRT2.1* was the most highly expressed member of the family among the 7 *NRT2*s (5 to 50 fold higher expression as compared to *NRT2.2, NRT2.4, NRT2.5*) (Supplemental Figure 2).

**Figure 2.**
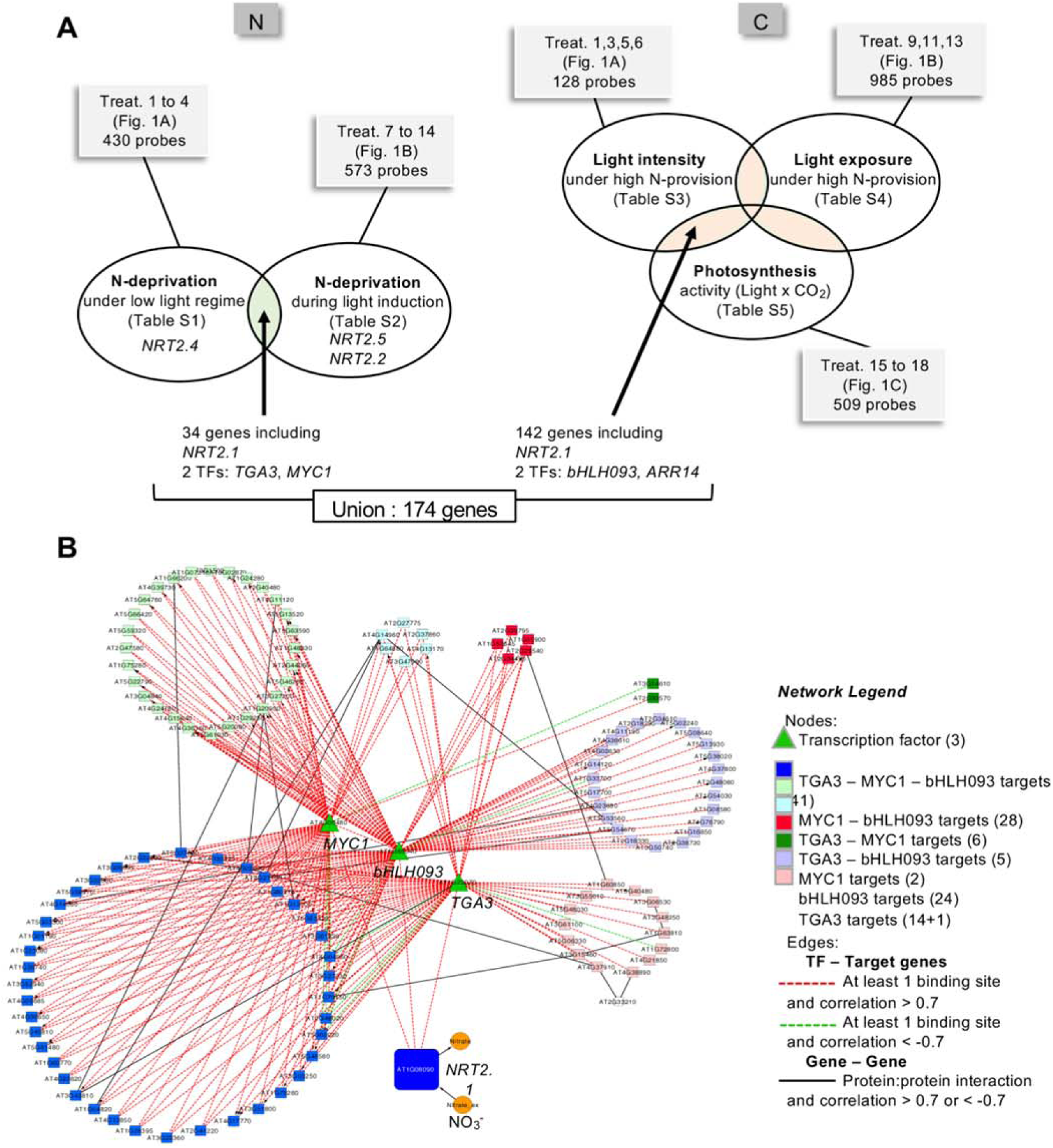
Gene expression multi-analysis driven by *NRT2.1* expression pattern combined to an integrative analysis identified a candidate gene regulatory network connected to the NO_3_^-^ transport system. **A.** Venn diagrams identifying common genes regulated by N provision on low light condition and dark to light transition (34 genes) or regulated by light/carbon (142 genes). The union of these gene lists defines a population of 174 genes, including 4 transcription factors. **B.** The core set of 174 genes differentially expressed has been structured into a Gene Regulatory Network using the Gene Networks analysis tool in VirtualPlant software (http://virtualplant.bio.nyu.edu/cgi-bin/vpweb/) *(32)*. The network includes 124 nodes (genes) and 260 edges connecting genes. The nodes have been organized according to their connection to the 3 transcription factors *MYC1, TGA3* and *bHLH093* and are detailed in the Network Legend. *ARR14* is excluded from the network due to its lack of connectivity to other nodes according to the edges selected to generate the network.

To find gene regulatory networks that could integrate N and C signalling and thus control *NRT2.1* expression, we defined 5 different subsets of conditions addressing the regulation by N on one side and by C on the other side, as described in Figure 2A. Genes defined as regulated by N-deprivation like *NRT2.1* are differentially regulated by N provision in conditions 1 to 4 in experiment 1, where *NRT2.4* is also found regulated and in conditions 7 to 14 in experiment 2, where *NRT2.2* and *NRT2.5* were also found regulated. To select the most robust genes regulated by N provision only the intersection between the 2 groups was isolated. In addition to *NRT2.1*, the intersection defines a set of 33 genes including the 2 transcription factors *TGA3* (At1g22070) and *MYC1* (At4g00480). On another hand, genes considered as regulated by C provision like *NRT2.1* are differentially regulated by light intensity in conditions 1, 3, 5, and 6 in experiment 1, by light time exposure in conditions 9, 11 and 13 in experiment 2 and by photosynthesis in conditions 15 to 18 in experiment 3. Similarly, to narrow down the specificity of gene regulation by C factor, only common genes to at least 2 experiments were isolated. This core set corresponds to 142 genes including *NRT2.1* but also 2 others transcription factors *bHLH093* (At5g65640) and *ARR14* (At2g01760) (Figure 2A).

Next, we only focused on the 174 genes that showed a response to N starvation (34 genes) and/or C provision (142 genes); *NRT2.1* being the common gene between the 2 responsive gene lists together with a *Kinesin3* gene (At5g54670-ATK3) coding for a microtubule motor protein. The possible connection of the 4 transcription factors with *NRT2.1* and the other genes was determined by a Gene Networks analysis performed on the VirtualPlant platform (Katari et al., 2010). The generated network contains 124 gene nodes. These genes are connected to each other by 260 edges, representing regulatory relationships such as predicted transcription factor-target gene interactions (Figure 2B). Regulatory interactions were proposed based on detection of at least one predicted binding site for a given transcription factor within the promoter region of the target gene as done previously (Gutierrez et al., 2008). According to the parameters used, 50 genes out of the 174 are not connected to any other genes in the network (See Material and Methods for details about the parameters). Among these 50 genes, the transcription factor *ARR14* was excluded due, for instance, to a low level of correlation between this gene and *NRT2.1* expression patterns. However, TGA3, MYC1 and bHLH093 have all predicted regulatory interactions with *NRT2.1* plus 40 other genes of the network (indicated in blue in Figure 2B). The network predicts also that only one or only two of these transcription factors putatively regulate the 79 remaining genes (one gene being connected to the network by predicted protein-protein interaction with 2 TGA3-targets). Nevertheless, almost all sub-networks are interconnected through protein-protein interaction prediction, suggesting possible coordination within the network at the whole.

### Regulation of *MYC1, TGA3* and *bHLH093* in response to C and N

The gene regulatory network we obtained revealed 3 main transcription factors: MYC1 and TGA3 which were found to be co-regulated with *NRT2.1* in response to N starvation and bHLH093 which was found to be co-regulated with *NRT2.1* in response to light/photosynthesis. In order to validate their regulation, we measured gene expression by QPCR across all the conditions performed in experiment 1 and 2 (Figure 3A). The results confirmed that expression of *TGA3* and *MYC1* genes is induced 2- to 3-fold after transferring the plants to a N-free solution, especially under LL or HL conditions. Furthermore, similar to *NRT2.1, MYC1* regulation of gene expression requires the presence of light (Figure 3A and Supplemental Figure 3). The results also confirmed that *bHLH093* gene expression is only induced by light (between 3- and 4-fold after 8h of HL), independent of N nutrition. This is supported by the fact that *bHLH093* is not regulated by N starvation (Figure 3A and Supplemental Figure 3). On the contrary, *MYC1* and *TGA3* genes are not only regulated by N starvation, but their expression is also induced by light, especially in plants starved for N (Figure 3A and Supplemental Figure 3). Like for *NRT2.1*, putative *cis*-binding elements for TGA3, MYC1 and bHLH093 were also found in the promoters of *NRT2.2, NRT2.4* and *NRT2.5* (Figure 3B). Furthermore, DNA affinity purification sequencing (DAP-seq) experiments recently performed by O’Malley et al. (2016) confirmed that TGA3 binds in silico to the promoter of *NRT2.1, NRT2.2* and *NRT2.4* (Figure 3C). Unfortunately, no data are available for MYC1 and bHLH093 in this work. Altogether, these results suggest that the transcription factors we identified are involved in regulation of several root *NRT2s*.

**Figure 3.**
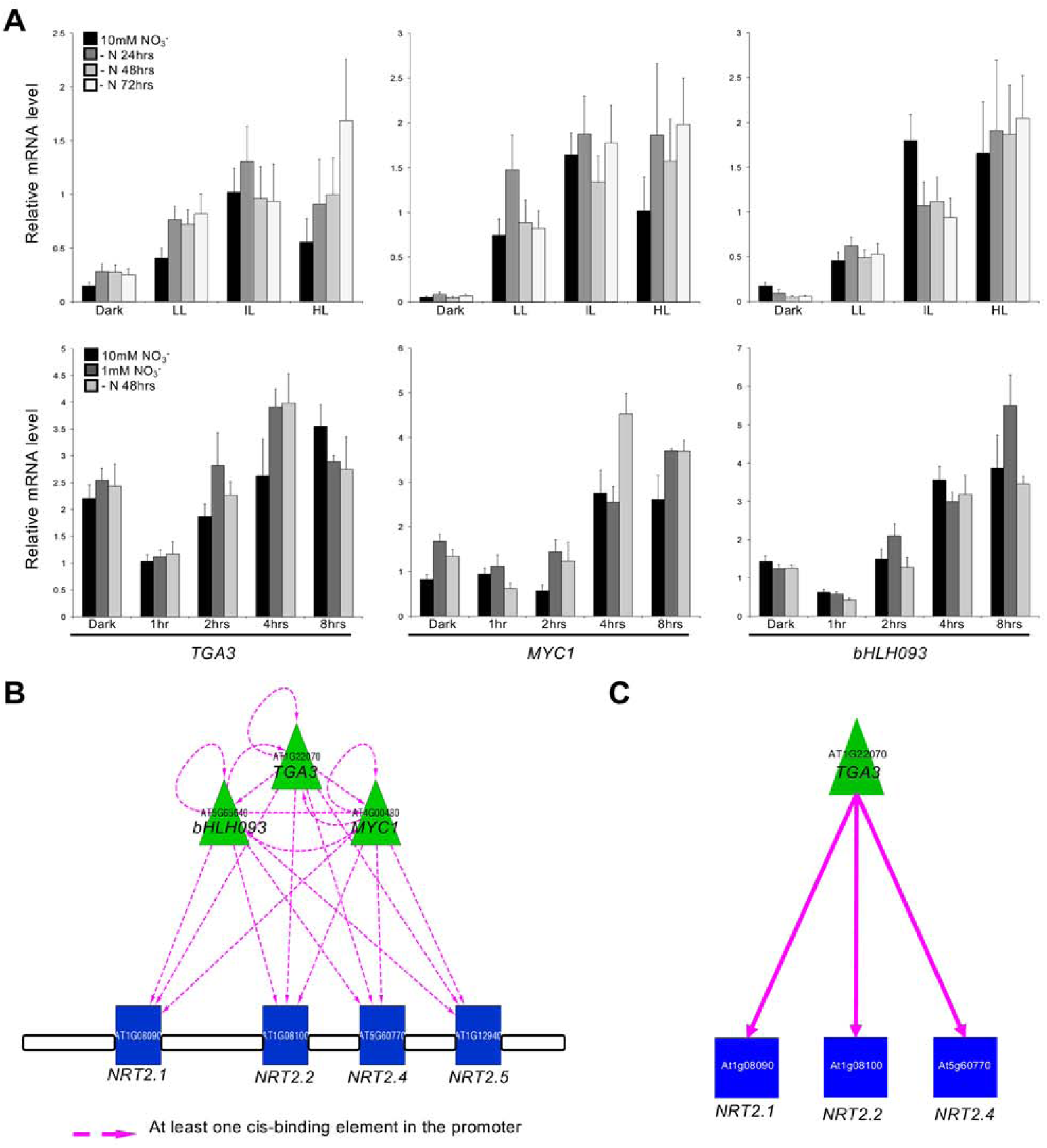
*TGA3, MYC1* and *bHLH093* are candidate transcription factors for the control of the expression of *NRT2* gene family. **A.** Gene expression analysis of the 3 candidate transcription factors in the extended set of Nitrogen/Carbon combinations confirms correlation with *NRT2.1* regulation. Expression patterns have been determined by RT-QPCR (relative accumulation to *Clathrin* housekeeping gene). **B.** *NRT2.2, NRT2.4* and *NRT2.5* as well as *NRT2.1* display putative cis-binding elements for the 3 transcription factors in their promoter region. The gene network has been done using the Gene Networks analysis tool in VirtualPlant software (http://virtualplant.bio.nyu.edu/cgi-bin/vpweb/) *(32)*; only Regulated Edges box and One Binding Site option has been selected in this case. **C.** TGA3 bounds *in silico* with the promoter of *NRT2.1, NRT2.2* and *NRT2.4*. The analysis has been done using the Plant Cistrome Database (http://neomorph.salk.edu/PlantCistromeDB) *(63)*.

To our knowledge, the transcription factors TGA3, MYC1 and bHLH093 have not been isolated in previous transcriptomic approach as candidates for regulation of root NO_3_^-^ transporters. In order to understand why they have not been found before we looked at the expression pattern of the known regulatory elements for *NRT2.1* in our experimental set up. The results show that the known regulators for *NRT2.1* were not co-regulated with *NRT2.1* expression in our conditions (Figure 4). This was also the case for *HY5*, a transcription factor recently identified as involved in the regulation of *NRT2.1* by light/photosynthesis (Chen et al., 2016). In our hands, this transcription factor was only induced by light independently of the presence of CO_2_ and therefore not by photosynthesis like *NRT2.1* (Figure 4). As most of the previous transcriptomic experiments were performed to study the signalling pathways involved in short-term induction by NO_3_^-^, we also looked at the regulation of *TGA3, MYC1* and *bHLH093* in those conditions (Supplemental Figure 4). We chose the transcriptomic experiments performed by Wang et al. (2004). In this study WT plants and the null mutant for nitrate reductase (NR) were treated with 5mM KNO_3_ for 2h and compared to control plants treated with 5mM KCl for 2h. The data sets allowed the authors to determine the genes that respond specifically to NO_3_^-^ in both WT and NR-null plants. The results show that, as expected, *NRT2.1, NRT2.2* and *NRT2.4* are induced by NO_3_^-^ while *NRT2.5* seems to be repressed (Supplemental Figure 4A). In the same time, most of the known regulators for *NRT2.1* are also induced by NO_3_^-^ except *NLP7* and *TCP20*, two transcription factors which have not been isolated using transcriptomic approaches (Supplemental Figure 4B). On the contrary, in the same conditions, our three transcription factors, *TGA3, MYC1* and *bHLH093* were not regulated by NO_3_^-^ supply neither in WT nor NR-null plants. All these results reinforced the originality of our experimental set up and explain why we found new candidates that have never been isolated in previous transcriptomic experiments.

**Figure 4.**
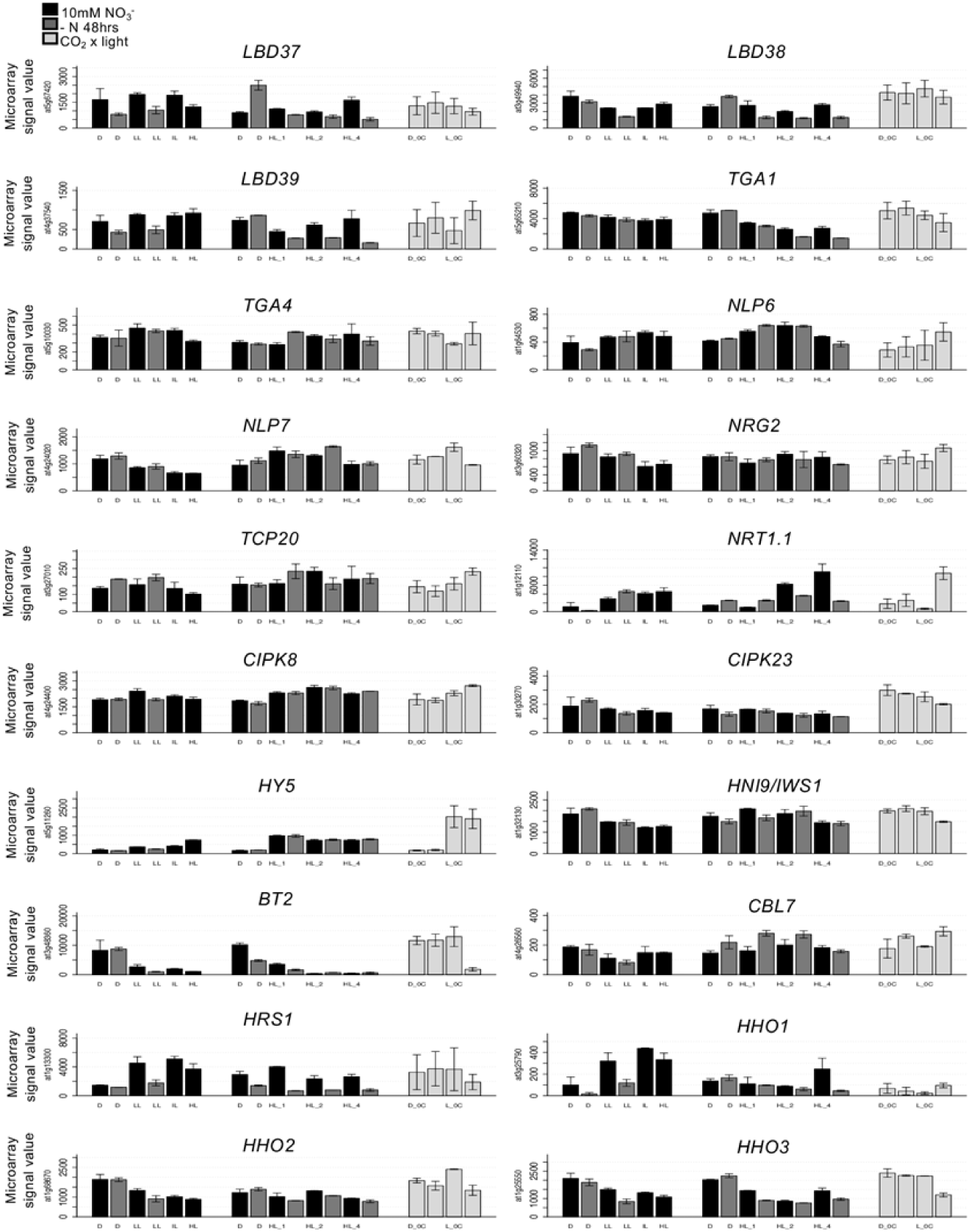
Most of the genes previously determined as *NRT2s* regulators do not display expression patterns similar to the patterns of the 3 candidate transcription factors in the set of Nitrogen and Light/Carbon combinations. Graphs display the expression pattern of the 20 genes extracted from the whole transcriptomic dataset. Data are organized according to the multi-analysis (*i.e.*, S1 to S5, Figure 2). LBD37, LBD38, LBD39 repress the expression of genes involved in NO_3_^-^ uptake (*NRT2.1* and *NRT2.5*) and assimilation, likely mimicking the effects of N organic compounds (Rubin et al., 2009). TGA1, TGA4, NLP6, NLP7, NRG2, NRT1.1, CIPK8, CIPK23 are required for the NO_3_^-^-dependent induction of *NRT2.1 (22, 23, 25, 27, 28, 45, 64, 65)*. TCP20 and HNI9/IWS1 are involved into *NRT2.1* regulation controlled by systemic signaling *(29, 66)*. BT2 represses expression of *NRT2.1* and *NRT2.4* genes under low NO_3_^-^ conditions *(26)*. CBL7 regulates *NRT2.4* and *NRT2.5* expression under N-starvation conditions *(40)*. HY5 has been recently identified as a regulator of *NRT2.1* by mediating light promotion of NO_3_^-^ uptake *(30)*. HRS1, HHO1, HHO2 and HHO3 are repressors of *NRT2.4* and *NRT2.5* expression under high N conditions *(41, 42).*

### Role of MYC1, TGA3 and bHLH093 in the regulation of NRT2s root nitrate transporters

To determine if MYC1, TGA3 and bHLH093 are involved in regulation of *NRT2* root NO_3_^-^ transporters we used two independent insertion mutants for each of the transcription factors: *tga3.2, tga3.3* for TGA3, *myc1.2, myc1.3* for MYC1 and *bHLH093.1, bHLH093.5* for bHLH093. As both *TGA3* and *MYC1* were found to be regulated by N starvation, we also produced a double mutant, *tga3.2*/*myc1.2*, to test a potential additive effect of those transcription factors on the regulation of *NRT2s*. In addition, to reinforce our conclusions concerning the role of bHLH093, we also produced an overexpressing line by transforming the *bhlh093.1* mutant with a *35S::bHLH093* construct. The measurement of *MYC1, TGA3* and *bHLH093* expression level confirmed an almost complete absence of their transcripts in their respective mutants and a strong overexpression of *bHLH093* in the overexpressing line (Supplemental Figure 5A and B).

**Figure 5.**
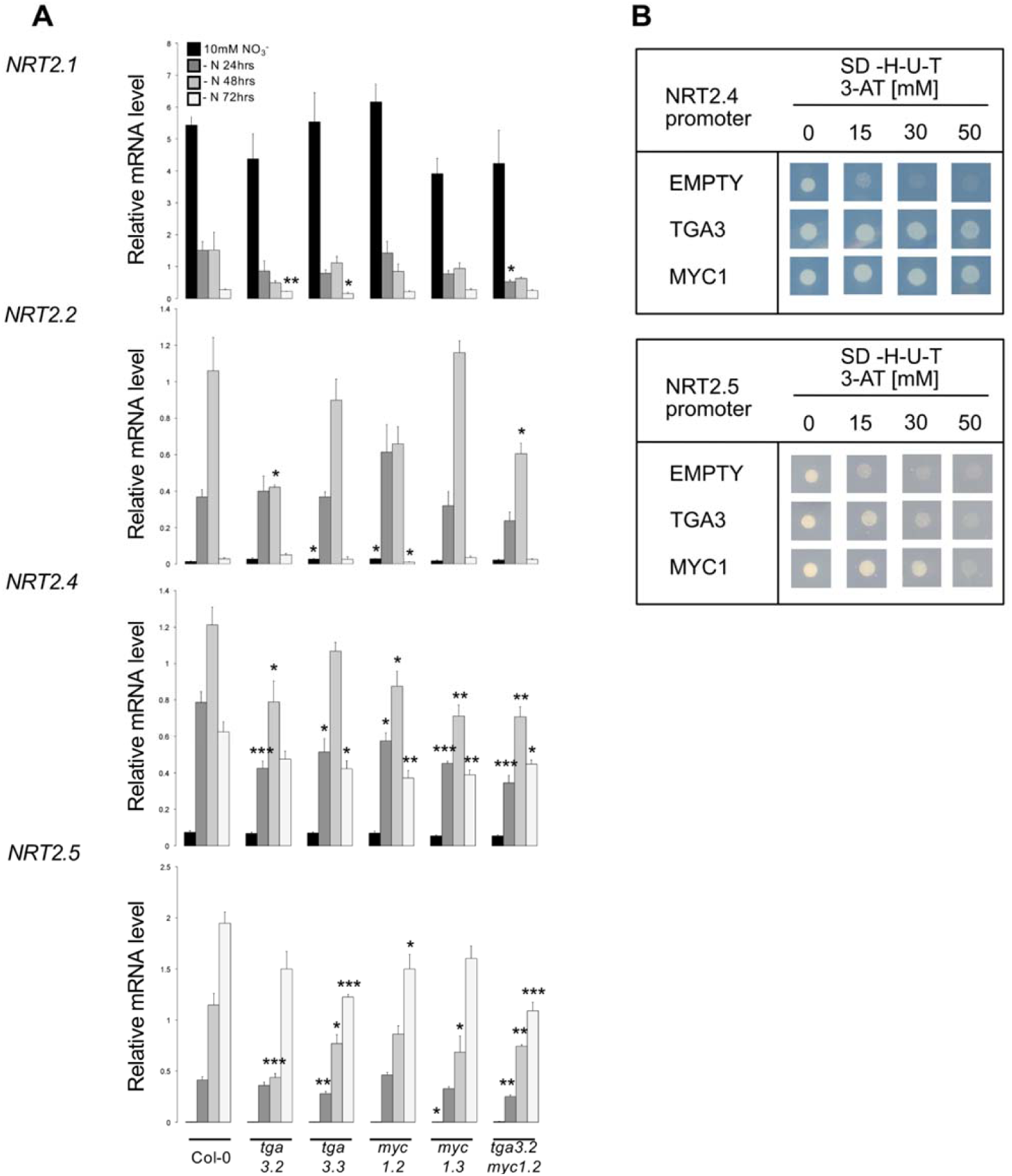
TGA3 and MYC1 are required for *NRT2.4* and *NRT2.5* full induction during N-deprivation. **A.** Characterization of the knock-out mutants for TGA3 (*tga3.2* and *tga3.3*), MYC1 (*myc1.2* and *myc1.3*) and the TGA3/MYC1 double mutants (*tga3.2 myc1.2*). The plants were supplied with NO ^-^ 10 mM one week ahead the experiment and acclimated for 24 hr in high light conditions (800 μmol m^-2^ s^-1^) before applying the N deprivation for 24, 48 or 72 hr. Roots have been collected to assess *NRT2.1, NRT2.2, NRT2.4* and *NRT2.5* mRNA accumulation by RT-QPCR (relative accumulation to *Clathrin* housekeeping gene). Values are means of three biological replicates ± SD. Differences between WT and the KO mutants are significant at * *P* < 0.05, * * *P* < 0.01, * * * *P* < 0.001 (Student’s *t* test). **B.** Characterization of TGA3 and MYC1 interaction with *NRT2.4* and *NRT2.5* promoters in a Y1H assay. Yeast cells were grown on SD-H-U-T minimal media without histidine (H), uracil (U), tryptophan (T) and containing 3- amino-1,2,4-triazole (3AT) at 0, 15, 30 and 50 mM. Interaction between the transcription factors and the promoters results in HIS3 reporter activation in contrast to the empty vector that does not interact.

As expected for a role of TGA3 and MYC1 in the regulation of *NRT2s* by N starvation, the induction of both *NRT2.4* and *NRT2.5* is overall reduced in *tga3* and *myc1* mutants compared to wild type plants, especially after 72h of N starvation (Figure 5A). This lower induction in response to N starvation is stronger in the double mutant *tga3.2*/*myc1.2* and is observed in that case consistently after 24h, 48h and 72h of N starvation for *NRT2.*4 and *NRT2.5* and after 48h for *NRT2.2*. It suggests that TGA3 and MYC1 are not redundant and that both factors may function as transcriptional activators under low N conditions. This result is supported by the fact that neither the level of expression nor the regulation of *MYC1* in *tga3* mutants and of *TGA3* in *myc1* mutants are affected compared to wild type plants (Supplemental Figure 5A). However, surprisingly, MYC1 and TGA3 do not affect the regulation of *NRT2.1* in the same conditions (Figure 5A). In agreement with a role of MYC1 and TGA3 in the regulation of *NRT2.4* and *NRT2.5*, Y1H experiments show that both transcription factors are able to bind to the promoter of these two transporters (Figure 5B).

Out of the three *NRT2s*, which are induced by light, *NRT2.4* and to a lower extent *NRT2.1*, have a significant lower induction after 4h and 8h of light in the *bHLH093* mutants as compared to wild type plants (Figure 6A and 6B). Conversely, the induction by light of both *NRT2.1* and *NRT2.4* is higher in the *35S::bHLH093* plants (Figure 6B). Interestingly, this phenotype seems to depend on the amount of NO_3_^-^ in the nutritive solution since the effect of bHLH093 is preferentially seen when plants are starved for N or on 1mM NO_3_^-^ and is almost absent when plants are grown on 10mM NO_3_^-^ (Figure 6A and 6B).

**Figure 6.**
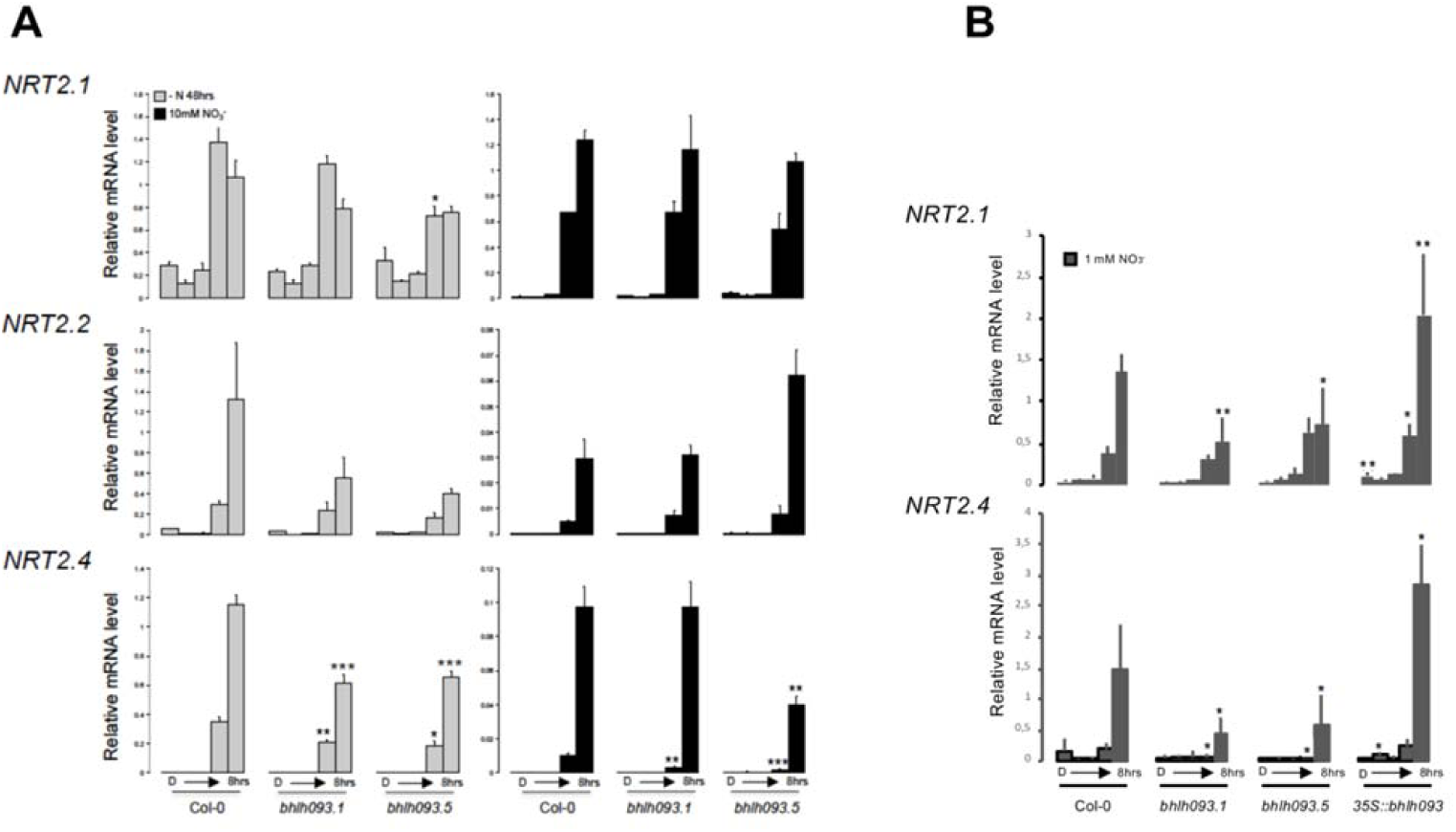
*bHLH093* is required for *NRT2.4* full induction by light in N-deprivation condition. **A.** Characterization of the knock-out mutants for bHLH093 (*bHLH093-1* and *bHLH093-5*) on 0N or 10mM NO_3_^-^. The plants were either starved for N for 48 hr (light gray bars) or supplied with NO_3_^-^ 10 mM one week ahead the experiment (black bars) and were kept in the dark 40 hr before transition to high light intensity (800 μmol m^-2^ s^-1^) during 1h, 2h, 4h and 8h. Roots have been collected to assess *NRT2.1, NRT2.2* and *NRT2.4* mRNA accumulation by RT-QPCR (relative accumulation to *Clathrin* housekeeping gene). Values are means of three biological replicates ± SD. Differences between WT (Col-0) and the KO mutants are significant at \**P* < 0.05, \*\**P* < 0.01, \*\*\**P* < 0.001 (Student’s *t* test). **B.** Characterization of the knock-out (*bHLH093-1* and *bHLH093-5*) and the over-expressor (*35S::bHLH093*) mutants for bHLH093 on 1mM NO_3_^-^. The plants were grown on 1mM NO_3_^-^ and were kept in the dark 40 hr before transition to intermediate light intensity (250 μmol m^-2^ s^-1^; IL) during 1h, 2h, 4h and 8h. Roots have been collected to assess *NRT2.1* and *NRT2.4* mRNA accumulation by RT-QPCR (relative accumulation to *Clathrin* housekeeping gene). Values are means of three biological replicates ± SD. Differences between WT (Col-0) and the mutants are significant at \**P* < 0.05, \*\**P* < 0.01, \*\*\**P* < 0.001 (Student’s *t* test).

## Discussion

### Interaction between nitrogen and light provision affect regulation of *NRT2.1* expression

As part of its central physiological role, the root NO_3_^-^ HATS is a main target of the C/N regulatory networks ensuring the necessary integration of both, N acquisition by roots and C acquisition by shoots. The HATS regulation by N starvation has been well characterised in previous studies, especially through the study of *NRT2.1* expression. Split-root experiments have demonstrated that this regulation relies on systemic signaling pathways (Gansel et al., 2001), and underlying molecular mechanisms have recently been unraveled (Ohkubo et al., 2017). On the other hand, *NRT2.1* expression is also dramatically induced by light and sugars through an Oxidative Pentose Phosphate Pathway (OPPP)-dependent signaling mechanism (Lejay et al., 1999; Lejay et al., 2003; Lejay et al., 2008; de Jong et al., 2014). Over the past decade, the importance of signal interaction for the regulation of gene expression has become more and more obvious and especially for C/N regulation (Palenchar et al., 2004; Gutierrez et al., 2007; Krouk et al., 2009). However, the details of how this interaction affects regulation of *NRT2.1* expression in response to combined N/C treatments were unknown. Our results clearly show that the interplay of N and C signaling mechanisms has a major role as light conditions can totally suppress N regulation of *NRT2.*1 expression, and vice versa (Figure 1A and 1B). Similar to the case for inorganic N assimilation, it seems that low sugars inhibit *NRT2.1* expression, overriding signals from N metabolism (Stitt et al., 2002; Nunes-Nesi et al., 2010). Surprisingly, the regulation of *NRT2.1* by N starvation is not only abolished when plants are treated in the dark. It happens also under high light conditions (Figure 1A). However, in that case, the level of *NRT2.1* expression is always high, even on normally repressive conditions like 10 mM NO_3_^-^, while in the dark the level of *NRT2.1* stays low, independently of the level of N. One model to explain these results is that enhancement of growth due to combination of high light and high NO_3_^-^ supply results in a sustained high N demand for growth, relieving the feedback repression normally associated with high NO_3_^-^ supply. This model is supported by a recent metabolomics analysis performed on *Arabidopsis thaliana* under diverse C and N nutrient conditions (Sato and Yanagisawa, 2014). Taken together, these results clearly support the idea that the control of *NRT2.1* expression involves a complex network of interactions between signals emanating from N and C metabolisms. However, this level of complexity seems to be rather specific for *NRT2.1*. In contrast to *NRT2.1*, expression of *NRT2.2, NRT2.4* and *NRT2.5* is always repressed on 10 mM NO_3_^-^, independent of light levels (Supplemental Figure 1A). It should be noted that in the N starvation experiments plants are transferred on a media with no N. This leads to the variation of two factors, the N status of the plants, which decreases when plants are starved for N, and the presence of NO_3_^-^ in the nutritive solution, which is suppressed by the transfer to N-free solution. Concerning the regulation of *NRT2.2, NRT2.4* and *NRT2.5* it is not known which one of these two factors is predominant since their expression was only measured in N starvation experiments (Kiba et al., 2012; Lezhneva et al., 2014; Kotur and Glass, 2015). It is thus possible that *NRT2.2, NRT2.4* and *NRT2.5* are only regulated locally by NO_3_^-^ and not by systemic signals of N demand. This idea is supported by the work of (Ma et al., 2015) showing that the regulation of *NRT2.4* and *NRT2.5* by N starvation depends on CBL7, which is specifically induced by NO_3_^-^ deficiency. Moreover, NIGT/HRS1s have been shown to act as transcriptional repressor of *NRT2.4* and *NRT2.5* upon NO_3_^-^ treatment (Kiba et al., 2018)Safi et al. 2018). Local regulation by NO_3_^-^ would explain why these transporters, unlike *NRT2.1*, are always repressed when plants are on 10 mM NO_3_^-^, regardless of the light conditions (Supplemental Figure 1A).

### Identification of three new candidates for regulation of *NRT2* genes using a systems biology approach

Over the past few years, transcriptomic approach and systems biology have been powerful tools to identify new regulatory elements involved in N signaling (For review (Medici and Krouk, 2014; Vidal et al., 2015). For root *NRT2s* genes and HATS activity in *Arabidopsis*, it enabled the identification of CIPK23 and CIPK8 in response to NO_3_^-^, LBDs transcription factors in response to high N and BT2, a negative regulator of *NRT2.1* and *NRT2.4* under low N conditions (Figure 7) (Ho et al., 2009; Hu et al., 2009; Rubin et al., 2009; Araus et al., 2016). For C and N signaling, previous microarray studies in response to transient treatments with NO_3_^-^, sucrose or NO_3_^-^ plus sucrose have been used to reveal, at the level of the genome, the existence of interaction between C and N signaling (Wang et al., 2003; Price et al., 2004; Scheible et al., 2004; Wang et al., 2004; Gutierrez et al., 2007; Huang et al., 2016). In Arabidopsis, over 300 genes have been found differentially expressed by combined C:N treatments compared to C or N treatments (Palenchar et al., 2004). However, because of the number of genes affected by C and/or N regulation and the complex interactions between the signalling pathways, none of these studies have led so far to the identification and the validation of new regulatory elements. The unexpected regulations of root *NRT2s* and especially of *NRT2.1* in our experimental set-up offer an interesting opportunity to find genes more specifically involved in the regulation of root NO_3_^-^ transporters by C and/or N, and to build a gene network model integrating regulators responding to N and/or C signals. As compared to previous transcriptomic approaches on N and C signaling in plants, we were able to narrow down the number of candidate genes by (i) using *NRT2.1* as a specific target and (ii) integrating the data from several Affymetrix microarrays to find gene networks coregulated with the expression of *NRT2.1* in response to different combinations of light and N treatments.

**Figure 7.**
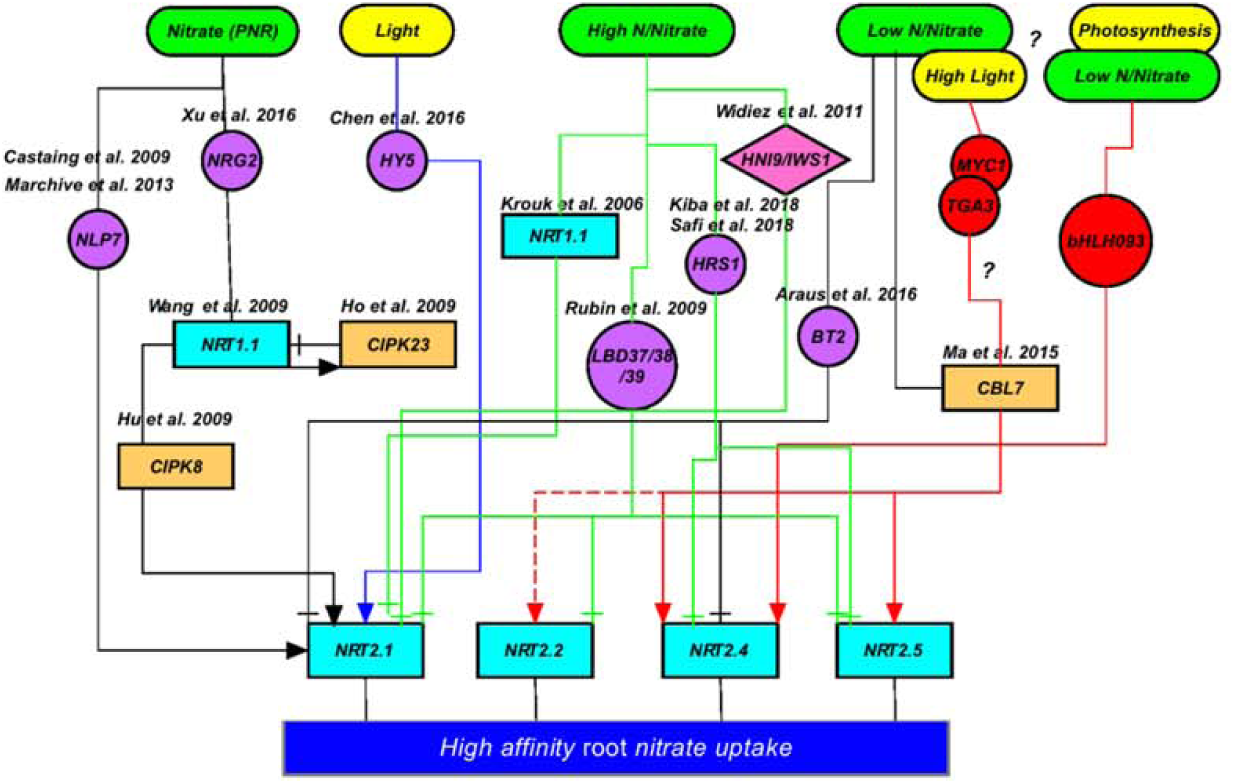
Schematic representation of the known regulatory elements for the regulation of root high-affinity NO_3_^-^ transporters in response to external NO_3_^-^, the N status of the plant and light/photosynthesis. Purple circles represent the transcription factors identified in previous studies while red circles represent the transcription factors identified in our study.

Therefore, the gene regulatory network includes only three transcription factors, bHLH093, MYC1 and TGA3 (Figure 2B). *bHLH093* was found co-regulated with *NRT2.1* in response to light through photosynthesis because, like *NRT2.1*, it is not induced by light in the absence of CO_2_ (Supplemental Figure 3). *MYC1* and *TGA3* were found co-regulated with *NRT2.1* in response to N starvation. The analysis of their level of expression across all the experiments revealed that *TGA3* and *MYC1* are induced by N starvation but especially in LL and HL conditions, while *bHLH093* seems overall induced by light no matter what the level of N (Figure 3A). Furthermore, *MYC1* is also clearly induced by light (Figure 3A and Supplemental Figure 3). Taken together these results support the validity of our approach to find regulatory elements affected by C and/or N signalling and which are thus candidates for the regulation by C/N interaction. Interestingly, none of these three transcription factors was found involved in the regulation of root NO_3_^-^ transporters by previous studies. One explanation to this relates to the fact that the expression of *bHLH093, MYC1* and *TGA3* is not responsive to the induction by NO_3_^-^, which was by far the major environmental change investigated by previous studies (Supplemental Figure 4A). Conversely, none of the regulatory genes identified in previous studies was found with our approach. Indeed, most of them are not affected by N starvation and/or by light (Figure 4). The only exception is *HY5*, which encodes a recently identified mobile transcription factor involved in the regulation of *NRT2.1* by sugar signals (Chen et al., 2016) and that is not found co-regulated with *NRT2.1* in our analysis. This is explained by the fact that, unlike *NRT2.1*, we found *HY5* induced by light even in the absence of CO_2_ in our dataset (Figure 4). It indicates that expression of HY5 does not depend of the production of sugars through photosynthesis and is directly regulated by light. The role of HY5 in light signalling and not in C signalling is supported by previous studies showing that HY5 works downstream phytochrome signalling (Quail, 2002; Li et al., 2010). Taken together, these results suggest that *NRT2.1* would be induced by both a light component dependent on HY5 and a C component dependent on the OPPP (Lejay et al., 2008; de Jong et al., 2014; Chen et al., 2016). Accordingly, both Lejay et al. (2008) and Chen et al. (2016) found that induction of *NRT2.1* by light is higher as compared to the addition of sucrose in the dark. Furthermore, there is still an induction of *NRT2.1* expression by increasing supply of sucrose in the mutant *hy5* (Chen et al., 2016).

### bHLH093, MYC1 and TGA3, three transcription factors involved in regulation of *NRT2.4* and *NRT2.5* gene expression

The use of mutants validated our approach and showed that bHLH093 has mainly a role in the induction by light of *NRT2.4*, while MYC1 and TGA3 affect induction by N starvation of both *NRT2.4* and *NRT2.5* and in a more modest way *NRT2.2* (Figure 5A and Figure 6). Furthermore, Y1H experiments support the fact that MYC1 and TGA3 are direct regulators of *NRT2.4* and *NRT2.5* as already suggested for TGA3 and *NRT2.4* by the results obtained by O’Malley et al. (2016) (Figure 3 and Figure 5B). Conversely, Chromatin Immunoprecipitation (ChIP) experiments, using plants expressing bHLH093 fused to GFP, failed to reveal a robust interaction with the promoter of *NRT2.4* (data not shown). It suggests that bHLH093 is an indirect regulator and that it is rather involved in the signalling pathway governing the regulation of *NRT2.4* and in a lesser extend *NRT2.1* by photosynthesis.

As represented in Figure 7, most of the regulatory elements identified to date concern the primary NO_3_^-^ response (PNR), with only three elements involved in the repression by high N or high NO_3_^-^ and one in the induction by light. Along with CBL7, MYC1 and TGA3 seem thus to be part of an independent signalling pathway involved in the induction of root NO_3_^-^ transporters in response to low N, while bHLH093 is, to our knowledge, the first element involved in a regulatory mechanism linked to photosynthesis (Ma et al., 2015). As discussed above, the role of these transcription factors in the regulatory mechanisms involved in C/N interactions is also supported by our results. Indeed the role of bHLH093 in the regulation by light seems to be dependent of the level of N and the role of MYC1 and TGA3 seems to be stronger in high light conditions (Figure 5A and Figure 6).

However, surprisingly, none of these 3 transcription factors affect strongly the regulation of *NRT2.1*, that we used as a target gene in our systems biology approach. This result could indicate that the regulatory mechanisms differ between the four *NRT2* genes involved in the HATS. Indeed, *NRT2.1* is regulated by at least 4 different mechanisms (local induction by NO_3_^-^ and repression by high NO_3_^-^, systemic repression by N metabolites and induction by C), while *NRT2.4* is regulated by C and N starvation and *NRT2.5* only by N starvation. Furthermore, our experimental setup revealed obvious complex interactions between N and C signalling for *NRT2.1,* which do not exist for *NRT2.2, NRT2.4* and *NRT2.5.* As discussed above, if *NRT2.2, NRT2.4* and *NRT2.5* are only repressed by NO_3_^-^ and not by N metabolites, MYC1 and TGA3 could be involved in a NO_3_^-^–specific signalling pathway upregulating the very high-affinity transporters (NRT2.4 and NRT2.5) when the external NO_3_^-^ concentration becomes too low to be efficiently taken up by NRT2.1. Previous results support a role, for at least TGA3, in a NO_3_^-^-specific signalling pathway. Indeed, TGA3 is part of a family of 7 genes in *Arabidopsis thaliana* and two of them, TGA1 and TGA4, have already been involved in the induction of *NRT2.1* and *NRT2.2* in response to NO_3_^-^ (Alvarez et al., 2014). Taken together these results and our findings suggest that this family of transcription factors could participate in a more general way to the regulation of root NO_3_^-^ transporters by NO_3_. Concerning MYC1 there is no direct evidence to support its role in a NO_3_ ^-^ signalling pathway (Bruex et al., 2012).

Since *NRT2.1* and *NRT2.4* are both regulated by C through OPPP, it was even more surprising to find that the absence of bHLH093 affects mainly the induction by light of *NRT2.4* compared to *NRT2.1* (Figure 6) (Lejay et al., 2008). However, the role of bHLH093 seems to be dependent on the level of N since it plays a significant role in the regulation of *NRT2.4* only under low N conditions, whereas the induction of *NRT2.*1 by light is mostly seen in this experiment under high N conditions (10mM NO_3_^-^). It could explain why, in those conditions, *bHLH093* mutation does not affect the regulation of *NRT2.1* by light, while in the second experiment, where plants were grown on a moderate level of NO_3_^-^ (1mM), *NRT2.1* is well induced by light and the NO_3_^-^ concentration could be low enough to reveal the impact of bHLH093 on *NRT2.1* regulation (Figure 6B). To our knowledge, the role of bHLH093 in the roots and in response to light has never been characterised before. The only information concerns a role in flowering promotion under non-inductive short-day conditions through the gibberellin pathway (Sharma et al., 2016).

## Materials and Methods

### Plant Material

*Arabidopsis thaliana* genotypes used in this study were the wild-type Col-0 ecotype and mutants obtained from the Salk Institute: *tga3.2* (Salk_081158), *tga3.3* (Salk_088114), *myc1.2* (Salk_057388), *myc1.3* (Salk_006354), *bHLH093.1* (Salk_121082) and *bHLH093.5* (Salk_104582).

In all experiments plants were grown hydroponically under non sterile conditions as described by Lejay et al. (1999). Briefly, the seeds were germinated directly on top of modified Eppendorf tubes filled with pre-wetted sand. The tubes were then positioned on floating rafts and transferred to tap water in a growth chamber under the following environmental conditions: light/dark cycle of 8 h/16 h, light intensity of 250 µmol·m^-2^·s^-1^, temperature of 22/20°C, and RH of 70%. After 1 week, the tap water was replaced with a complete nutrient solution. The experiments were performed on plants grown on 1 mM NO_3_^-^ as N source. The other nutrients were added as described by Lejay et al. (1999). The plants were allowed to grow for 3 additional weeks before the experiments. Nutrient solutions were renewed weekly and on the day before the experiments.

### Treatments

Two different sets of experiments were performed to (i) study the impact of light on the regulation of NO_3_^-^ transporter genes in the roots by N starvation, and (ii) study the impact of the N status of the plants on the regulation of these genes by light.

In the first set of experiments 4 weeks old plants were transferred on a solution containing 10 mM NO_3_^-^. After one week the plants were transferred in the morning either in continuous dark or in a light/dark cycle at three different light intensities (50, 250 and 800 μmoles.h^-1^.m^-2^) and starved for N during 24h, 48h and 72h, by replacing NO_3_^-^ with CaCl_2_ 2.5 mM and K_2_SO_4_ 2.5 mM.

In the second set of experiments 4 weeks old plants were transferred on a solution containing 10 mM NO_3_^-^. They were then pre-treated during 3 days on nutrient solution containing contrasted level of N: (i) no N, (ii) 1 mM NO_3_^-^ or (iii) 10 mM NO_3_^-^. After 32h in the dark the plants were transferred to light for 1h, 2h, 4h and 8h under three different light intensities (50, 250 and 800 μmoles.h^-1^.m^-2^).

The dependence of the expression of NO_3_^-^ transporter genes on photosynthesis was investigated by modifying the CO_2_ concentration in the atmosphere. After a pretreatment of 40 h in the dark, plants grown on 1mM NO_3_^-^ were placed for 4 h in the light (□150 µmol·m^-2^·s^-1^) or in the dark in a 240-L, airtight plexiglass chamber connected to a computerized device for controlling temperature, humidity, and CO_2_ concentration in the atmosphere (Atelliance Instruments; see Delhon et al. (1996) for details). The CO_2_ concentration in the atmosphere was held constant during the treatments at 0 or 600 *μ*L L^-1^. All experiments were repeated two or three times.

### RNA Extraction and Gene Expression Analysis

Root samples were frozen in liquid N_2_ in 2-mL tubes containing one steel bead (2.5 mm diameter). Tissues were disrupted for 1 min at 30 s^-1^ in a Retsch mixer mill MM301 homogenizer (Retsch, Haan, Germany). Total RNA was extracted from tissues using TRIzol reagent (Invitrogen, Carlsbad, CA, USA). Subsequently 4 µg of RNA were treated with DNase (DNase I, SIGMA-ALDRICH, USA) following the manufacturer’s instructions. Reverse transcription was achieved in the presence of Moloney murine leukemia virus reverse transcriptase (Promega, Madison, WI, USA) after annealing with an anchored oligo(dT)_18_primer as described by Wirth et al. (2007). The quality of the cDNA was verified by PCR using specific primers spanning an intron in the gene *APTR* (At1g27450) forward 5’- CGCTTCTTCTCGACACTGAG-3’; reverse 5’-CAGGTAGCTTCTTGGGCTTC-3’.

Gene expression was determined by quantitative real-time PCR (LightCycler; Roche Diagnostics, Basel, Switzerland) with the kit LightCycler FastStart DNA Master SYBR Green I (Roche Diagnostics, Basel, Switzerland) according to the manufacturer’s instructions with 1 µl of cDNA in a total volume of 10 µl. The amplifications were performed as described previously by Wirth et al. (2007). All the results presented were standardized using the housekeeping gene Clathrin (At4g24550). Gene-specific primer sequences were: NRT2.1 forward, 5’-AACAAGGGCTAACGTGGATG-3’; NRT2.1 reverse, 5’- CTGCTTCTCCTGCTCATTCC-3’; NRT2.2 forward, 5’-GCAGCAGATTGGCATGCATTT- 3’; NRT2.2 reverse, 5’-AAGCATTGTTGGTTGCGTTCC-3’; NRT2.4 forward, 5’-GAACAAGGGCTGACATGGAT-3’; NRT2.4 reverse, 5’-GCTTCTCGGTCTCTGTCCAC-3’; NRT2.5 forward, 5’-TGTGGACCCTCTTCCAAAAA-3’; NRT2.5 reverse, 5’- TTTGGGGATGAGTCGTTGTGG-3’; MYC1 forward, 5’-AACCTTAACGACTCTGTG-3’; MYC1 reverse, 5’-CCGCAACTATGTAGTCTCTG-3’; TGA3 forward, 5’- CTCTCAGAAAGTGTTGGC-3’; TGA3 reverse, 5’-CATATACGAGGAGATGAGTG-3’; bHLH093 forward, 5’-AGCTTGAAGGCCAACC-3’; bHLH093 reverse, 5’-GCTCTTTCATGTAATCTATGGCA-3’; Clathrin forward, 5’- AGCATACACTGCGTGCAAAG-3’; Clathrin reverse, 5’-TCGCCTGTGTCACATATCTC- 3’.

### Acquisition of Genome-Wide Expression and Statistical Analysis

Genome-wide expression was determined using Affymetrix ATH1 GeneChip expression microarrays according to manufacturer’s instructions. To do so, biotinylated cRNA was synthesized from 200 ng of total RNA from *Arabidopsis* roots. Affymetrix data were normalized in R (http://www.r-project.org/) using MAS5.

Then, normalized data were subjected to different statistical analyses, all centered on *NRT2.1* expression pattern but including various sets of microarray data among the whole data set. As a first approach to build a gene network involved in the regulation of root NO_3_^-^ transporters, we examined genes displaying expression pattern correlated to *NRT2.1* expression pattern across the entire dataset. A R^2^ coefficient cut-off above 0.8 or below −0.8 led to the identification of 79 AGIs displaying an expression pattern correlated to *NRT2.1*, including 77 genes positively correlated with *NRT2.1* (Table S1). Among these 79 genes, none of them displays a function related to gene regulation but rather related to metabolic activity and more precisely to carboxylic acid metabolic process as, for example, the *Glutamate synthase 2* gene (Supplemental Figure 6A). Moreover, a hierarchical clustering of the treatments according to the expression pattern of these genes clearly revealed that their response is largely driven only by the light/carbon factor, putting aside any possible regulation by N provision (Supplemental Figure 6B). Therefore, we determined that a global analysis of the entire data set was not relevant to identify regulators of NO_3_^-^ transport integrating C and N availability and that a finest analysis of gene expression in different subsets of treatments will be more powerful. The list of genes regulated by N-deprivation specifically under low light regime was determined by a t.test analysis (p.value<0.05) between conditions 3 and 4. All genes also found regulated between conditions 1 and 2 based on the same analysis are removed from this list (Figure 1, Table S1). Genes regulated by N-deprivation during light induction are determined by a 2 ways ANOVA using Nitrogen as one factor (presence = conditions 7,9,11,13 / absence = conditions 8,10,12,14) and Light as the second factor (no Light = conditions 7,8 / 1hr-light = conditions 9,10 / 2hr-light = conditions 11,12 / 4hr-light = conditions 13,14). Genes of interest are regulated by the interaction of the 2 factors (p.value<0.05) and display a similar regulation by N from dark to 2hr-light as observed for *NRT2.1* (Figure 1, Table S2). Genes regulated by light intensity under high N-provision and by light time exposure under high N-provision are both determined by a linear modeling of gene expression across light intensity (conditions 1,3,5,6) or time exposure (conditions 7,9,11,13) using a R^2^ above 0.9 (p.value is below 0.003) (Figure 1, Tables S3 and S4). Finally, genes regulated by photosynthesis activity are determined by a 2 ways ANOVA using CO_2_ level as one factor (0ppm = conditions 15,17 / 600ppm = conditions 16,18) and Light as the second factor (Dark = conditions 15,16 / Light = conditions 17,18). To narrow down the list of *NRT2.1*-like genes, only those passing post-hoc Tukey tests comparing conditions 18 to all 3 others (p.value<0.05) and displaying a ratio >2 or <0.5 are selected (Figure 1, Table S5).

### Visualization of gene connectivity by clustering and gene network analysis

Heat map hierarchical cluster of gene expression and samples was generated with the MeV software using Pearson correlation as distance metric and Average as linkage method (www.tm4.org) (Saeed et al., 2003). The Gene Network was generated with the VirtualPlant 1.3 software (http://virtualplant.bio.nyu.edu/cgi-bin/vpweb/) (Katari et al., 2010). The connectivity of the nodes is based on 5 categories corresponding to literature data, post-transcriptional regulation, protein:protein interactions, transcriptional regulation and regulated edges meaning transcription factor - target relationship based at least on one binding site in the promoter of the target gene. Two nodes are linked by an edge if they fall in any of these categories combined to an expression pattern correlated at a R^2^>0.7 or <-0.7. Visualization of the gene regulatory network has been performed with Cytoscape (http://www.cytoscape.org/) (Shannon et al., 2003). Node properties have been modified to reveal connectivity with the 3 transcription factors and highlight *NRT2.1* position within the network.

### Y1H Assays

For the generations of the plasmids for promoter analysis by Y1H, particular promoter fragments of NRT2.4 (1968bp), NRT2.5 (1692bp) were first amplified by PCR with overlapping ends as described by Gibson et al. (2009). For the bait, the pMW2 and pMW3 vectors were used (Deplancke et al., 2006). pMW vectors were amplified by PCR with overlapping ends as a single sequence (pMW2) or as 2 independent sections (pMW2). Final vectors were made as described by Gibson et al., 2009. The Y1H prey vectors for TGA3 and MYC1 transcriptions factors were a kind gift from Franziska Turck (Castrillo et al., 2011). All the fragments generated for all constructs were validated by DNA sequencing.

The Y1H assay was performed according to protocol described by Grefen (2014) with minor modifications. Briefly, the vectors pMW2-NRT2.4, pMW3-NRT2.4, pMW2-NRT2.5, pMW3-NRT2.5 were first linearized with restriction enzymes. For pMW2 vectors BamH1 (NEB) was used and for pMW3 vectors Xho1 (NEB). The resulting linearized constructs were subsequently co-integrated into the yeast strain: YM4271 as described by Grefen (2014). The transformed yeast strains were tested for autoactivation and the selected colonies with the higher sensitivity to 3-AT were then transformed with the construct pDEST-AD-TGA3 or pDEST-AD-MYC1 or pDEST-AD (Empty vector). Empty vector was included as a negative control. Resulting yeast were dropped on selection media (SD –His–Ura–Trp) supplemented with increasing concentrations of 3-AT (0, 15, 30, 50, 80, 100 mM). Yeast growth was verified after 48h.

## Acknowledgments

We thank members of the lab in France and Chile for discussion.

---

**Figure 1**. Interaction between Nitrogen and Light/Carbon provision modulates *NRT2.1* mRNA accumulation in roots. A, Different light regimes modulate *NRT2* regulation in roots of plants experiencing from high NO_3_^-^ provision (10 mM) to N deprivation (-N). The light regimes encompass dark, low light intensity (50 μmol m^-2^ s^-1^; LL), intermediate light intensity (250 μmol m^-2^ s^-1^; IL) and high light intensity (800 μmol m^-2^ s^-1^; HL). Plants were supplied with NO_3_^-^ 10 mM one week ahead the experiment and acclimated for 24 hours in the different light regimes before applying the N deprivation for 24, 48 or 72 hr. B, Different N provisions modulate *NRT2* regulation in roots of plants experiencing a dark to light transition. The N provisions encompass 10mM NO_3_^-^, 1mM NO_3_^-^ (for 72 hr) and N deprivation for 48 hr (-N). Plants are kept in the dark (*i.e.*, 40hr) before transition to high light intensity (800 μmol m^-2^ s^-1^; HL) and roots are collected at time 0 (Dark) and 1, 2, 4 and 8 hr after light transition. C, Regulation of *NRT2* by photosynthesis activity. Plants are grown in regular NO_3_^-^ regime (1mM) and intermediate light intensity until they are transferred for 4 hr in a CO_2_-deprived atmosphere (0ppm) or in high CO_2_-supplied atmosphere (600ppm), either in the dark or in the light. In these 3 experimental conditions, roots have been collected to assess *NRT2.1* mRNA accumulation by RT-QPCR (relative accumulation to *Clathrin* housekeeping gene). Expression pattern of *NRT2.1* across the 35 conditions tested (16 in A, 15 in B and 4 in C) has driven the choice of 18 conditions to investigate gene reprogramming associated to the regulation of NO_3_^-^ transport. These 18 conditions are indicated with arrows and numbers on the x-axis of the 3 *NRT2.1* bar graphs (Each arrow corresponds to one condition with 2 independent biological repeats constituted of a pool of approx. 10 plants each).

**Figure 2**. Gene expression multi-analysis driven by *NRT2.1* expression pattern combined to an integrative analysis identified a candidate gene regulatory network connected to the NO_3_^-^ transport system. A, Venn diagrams identifying common genes regulated by N provision on low light condition and dark to light transition (34 genes) or regulated by light/carbon (142 genes). The union of these gene lists defines a population of 174 genes, including 4 transcription factors. B, The core set of 174 genes differentially expressed has been structured into a Gene Regulatory Network using the Gene Networks analysis tool in VirtualPlant software (http://virtualplant.bio.nyu.edu/cgi-bin/vpweb/) (Katari et al., 2010). The network includes 124 nodes (genes) and 260 edges connecting genes. The nodes have been organized according to their connection to the 3 transcription factors *MYC1, TGA3* and *bHLH093* and are detailed in the Network Legend. *ARR14* is excluded from the network due to its lack of connectivity to other nodes according to the edges selected to generate the network.

**Figure 3.** *TGA3, MYC1* and *bHLH093* are candidate transcription factors for the control of the expression of *NRT2* gene family. A, Gene expression analysis of the 3 candidate transcription factors in the extended set of Nitrogen/Carbon combinations confirms correlation with *NRT2.1* regulation. Expression patterns have been determined by RT-QPCR (relative accumulation to *Clathrin* housekeeping gene). B, *NRT2.2, NRT2.4* and *NRT2.5* as well as *NRT2.1* display putative cis-binding elements for the 3 transcription factors in their promoter region. The gene network has been done using the Gene Networks analysis tool in VirtualPlant software (http://virtualplant.bio.nyu.edu/cgi-bin/vpweb/) (Katari et al., 2010); only Regulated Edges box and One Binding Site option has been selected in this case. C, TGA3 bounds *in silico* with the promoter of *NRT2.1, NRT2.2* and *NRT2.4*. The analysis has been done using the Plant Cistrome Database (http://neomorph.salk.edu/PlantCistromeDB) (O’Malley et al., 2016).

**Figure 4.** Most of the genes previously determined as *NRT2s* regulators do not display expression patterns similar to the patterns of the 3 candidate transcription factors in the set of Nitrogen and Light/Carbon combinations. Graphs display the expression pattern of the 20 genes extracted from the whole transcriptomic dataset. Data are organized according to the multi-analysis (*i.e.*, S1 to S5, Figure 2). LBD37, LBD38, LBD39 repress the expression of genes involved in NO_3_^-^ uptake (*NRT2.1* and *NRT2.5*) and assimilation, likely mimicking the effects of N organic compounds (Rubin et al., 2009). TGA1, TGA4, NLP6, NLP7, NRG2, NRT1.1, CIPK8, CIPK23 are required for the NO_3_^-^-dependent induction of *NRT2.1* (Munos et al., 2004; Castaings et al., 2009; Ho et al., 2009; Hu et al., 2009; Konishi and Yanagisawa, 2013; Marchive et al., 2013; Alvarez et al., 2014; Xu et al., 2016). TCP20 and HNI9/IWS1 are involved into *NRT2.1* regulation controlled by systemic signaling (Widiez et al., 2011; Guan et al., 2014). BT2 represses expression of *NRT2.1* and *NRT2.4* genes under low NO_3_^-^ conditions (Araus et al., 2016). CBL7 regulates *NRT2.4* and *NRT2.5* expression under N-starvation conditions (Ma et al., 2015). HY5 has been recently identified as a regulator of *NRT2.1* by mediating light promotion of NO_3_^-^ uptake (Chen et al., 2016). HRS1, HHO1, HHO2 and HHO3 are repressors of *NRT2.4* and *NRT2.5* expression under high N conditions (Kiba et al., 2018)Safi et al. 2018)

**Figure 5.** TGA3 and MYC1 are required for *NRT2.4* and *NRT2.5* full induction during N-deprivation. A, Characterization of the knock-out mutants for TGA3 (*tga3.2* and *tga3.3*), MYC1 (*myc1.2* and *myc1.3*) and the TGA3/MYC1 double mutants (*tga3.2 myc1.2*). The plants were supplied with NO_3_^-^ 10 mM one week ahead the experiment and acclimated for 24 hr in high light conditions (800 μmol m^-2^ s^-1^) before applying the N deprivation for 24, 48 or 72 hr. Roots have been collected to assess *NRT2.1, NRT2.2, NRT2.4* and *NRT2.5* mRNA accumulation by RT-QPCR (relative accumulation to *Clathrin* housekeeping gene). Values are means of three biological replicates ± SD. Differences between WT and the KO mutants are significant at \**P* < 0.05, \*\**P* < 0.01, \*\*\**P* < 0.001 (Student’s *t* test). B, Characterization of TGA3 and MYC1 interaction with *NRT2.4* and *NRT2.5* promoters in a Y1H assay. Yeast cells were grown on SD-H-U-T minimal media without histidine (H), uracil (U), tryptophan (T) and containing 3- amino-1,2,4-triazole (3AT) at 0, 15, 30 and 50 mM. Interaction between the transcription factors and the promoters results in HIS3 reporter activation in contrast to the empty vector that does not interact.

**Figure 6.** *bHLH093* is required for *NRT2.4* full induction by light in N-deprivation condition. A, Characterization of the knock-out mutants for bHLH093 (*bHLH093-1* and *bHLH093-5*) on 0N or 10mM NO_3_^-^. The plants were either starved for N for 48 hr (light gray bars) or supplied with NO_3_^-^ 10 mM one week ahead the experiment (black bars) and were kept in the dark 40 hr before transition to high light intensity (800 μmol m^-2^ s^-1^) during 1h, 2h, 4h and 8h. Roots have been collected to assess *NRT2.1, NRT2.2* and *NRT2.4* mRNA accumulation by RT-QPCR (relative accumulation to *Clathrin* housekeeping gene). Values are means of three biological replicates ± SD. Differences between WT (Col-0) and the KO mutants are significant at \**P* < 0.05, \*\**P* < 0.01, \*\*\**P* < 0.001 (Student’s *t* test). B, Characterization of the knock-out (*bHLH093-1* and *bHLH093-5*) and the over-expressor (*35S::bHLH093*) mutants for bHLH093 on 1mM NO_3_^-^. The plants were grown on 1mM NO_3_^-^ and were kept in the dark 40 hr before transition to intermediate light intensity (250 μmol m^-2^ s^-1^; IL) during 1h, 2h, 4h and 8h. Roots have been collected to assess *NRT2.1* and *NRT2.4* mRNA accumulation by RT-QPCR (relative accumulation to *Clathrin* housekeeping gene). Values are means of three biological replicates ± SD. Differences between WT (Col-0) and the mutants are significant at \**P* < 0.05, \*\**P* < 0.01, \*\*\**P* < 0.001 (Student’s *t* test).

**Figure 7.** Schematic representation of the known regulatory elements for the regulation of root high-affinity NO_3_^-^ transporters in response to external NO_3_^-^, the N status of the plant and light/photosynthesis. Purple circles represent the transcription factors identified in previous studies while red circles represent the transcription factors identified in our study.

**Supplemental Figure 1**. Interaction between Nitrogen and Light/Carbon provision modulates mRNA accumulation in roots of most of the *NRT2* family members.

**Supplemental Figure 2**. Expression pattern of *NRT2* family genes in the set of Nitrogen and Carbon/Light combinations as determined by Arabidopsis Affymetrix ATH1 microarray hybridization.

**Supplemental Figure 3**. Expression pattern of *TGA3, MYC1* and *bHLH093* transcription factors in the set of Nitrogen and Carbon/Light combination as determined by *Arabidopsis* Affymetrix ATH1 microarrays hybridization.

**Supplemental Figure 4**. *TGA3, MYC1* and *bHLH093* display expression pattern different than most of the known regulators of *NRT2* genes in response to NO_3_^-^.

**Supplemental Figure 5.** Expression pattern of *TGA3, MYC1* and *bHLH093*.

**Supplemental Figure 6.** Biomaps and hierarchical clustering of the 79 most correlated genes to *NRT2.1* expression across all experiments.

**Supplemental Table 1.** List of 430 probes regulated by N deprivation under low light regime only.

**Supplemental Table 2.** List of 573 probes regulated by the interaction between nitrogen and light.

**Supplemental Table 3.** List of 128 probes linearly regulated by light intensity.

**Supplemental Table 4.** List of 985 probes linearly regulated during light induction.

**Supplemental Table 5.** List of 509 probes regulated by the interaction between light and CO_2_.

**Supplemental Table 6.** List of 80 probes coregulated based on *NRT2.1* expression and pearson correlation.

